# BEEtag : a low-cost, image-based tracking system for the study of animal behavior and locomotion

**DOI:** 10.1101/020347

**Authors:** James D Crall, Nick Gravish, Andrew M. Mountcastle, Stacey A. Combes

## Abstract

A fundamental challenge common to studies of animal movement, behavior, and ecology is the collection of high-quality datasets on spatial positions of animals as they change through space and time. Recent innovations in tracking technology have allowed researchers to collect large and highly accurate datasets on animal spatiotemporal position while vastly decreasing the time and cost of collecting such data. One technique that is of particular relevance to the study of behavioral ecology involves tracking visual tags that can be uniquely identified in separate images or movie frames. These tags can be located within images that are visually complex, making them particularly well suited for longitudinal studies of animal behavior and movement in naturalistic environments. While several software packages have been developed that use computer vision to identify visual tags, these software packages are either (a) not optimized for identification of single tags, which is generally of the most interest for biologist, or (b) suffer from licensing issues, and therefore their use in the study of animal behavior has been limited. Here, we present BEEtag, an open-source, image-based tracking system in Matlab that allows for unique identification of individual animals or anatomical markers. The primary advantages of this system are that it (a) independently identifies animals or marked points in each frame of a video, limiting error propagation, (b) performs well in images with complex background, and (c) is low-cost. To validate the use of this tracking system in animal behavior, we mark and track individual bumblebees (*Bombus impatiens*) and recover individual patterns of space use and activity within the hive. Finally, we discuss the advantages and limitations of this software package and its application to the study of animal movement, behavior, and ecology.

## Introduction

A fundamental challenge facing diverse fields of research is the accurate reconstruction of spatial position information over time. In biology, for example, fields such as biomechanics, animal behavior, and ecology all depend heavily on reconstructing accurate spatio-temporal data on either anatomical components (e.g. different joints) of animals or their entire bodies. Traditionally, such tracking has been done primarily through human observation or manual tracking of positional information. Studies of animal locomotion, for example, have traditionally involved manual (although often computer-aided) tracking of anatomical features to reconstruct accurate movement kinematics (Tobalske et al., 2007; Wakeling and Ellington, 1997). On the other hand, studies of animal behavior and ecology have often involved marking animals with uniquely identifiable tags combined with manual observation (Seeley et al., 1991).

While such data sets have been indispensable for advancing their respective fields, manual collection of these data is time-intensive, laborious, and poorly-suited to generating large datasets, particularly those that involve tracking either multiple individuals or body parts simultaneously. In recent decades, advances in tracking technology have allowed researchers to collect large, highly accurate datasets in a fraction of the time taken by manual methods. For example, semi-automated marker tracking (Hedrick, 2008) or visual hull reconstruction (Ristroph et al., 2009) have allowed for the collection of highly accurate spatio-temporal datasets on animal locomotion. In ethology, automated tracking techniques have allowed for the collection of vast, highly-accurate behavioral datasets (Dell et al., 2014; Kain et al., 2013; Pérez-Escudero et al., 2014), which can be used, for example, in detailed quantitative analysis of animal behavior (Berman et al., 2014; Mersch et al., 2013).

A fundamental limit of many of the tracking methods described above, however, is the need for a controlled, laboratory environment for high-quality tracking results, which for certain research questions can present a significant limitation. Partially for this reason, radio-frequency identification (RFID) technology, which does not require a controlled visual environment for identification, has become particularly popular among behavioral ecologists for tracking and identifying individuals in both vertebrate (see Bonter and Bridge, 2011 for an excellent review of the use of this technology in birds) and invertebrate (Henry et al., 2012; Stelzer and Chittka, 2010) animals. While robust to limitations of the visual environment, however, the spatial information provided by RFID is limited, since spatial position is only recorded when an animal is near an RFID reader, and the technology is therefore of limited utility for addressing certain experimental questions.

Increasingly, automated image-based tracking has been used to explore basic questions in behavior and ecology (Dell et al., 2014). However, each tracking method has distinct strengths and limitations. One limitation that faces many image-based individual tracking methods is error propagation: since tracking is often based on using information from previous frames in a movie (e.g. spatial proximity of an animal from one frame to the next (Branson et al., 2009; de Chaumont et al., 2012; Hedrick, 2008)), errors can be introduced when the paths of two animals cross. Such errors are generally irreversible and propagate through time, thus making it difficult or impossible to track individuals over long time periods. While computational advances can reduce (Branson et al., 2009) or nearly eliminate (Pérez-Escudero et al., 2014) this problem, these techniques still rely on controlled, homogenous visual environments for accurate tracking.

One method for avoiding such errors and allowing for long-term tracking of uniquely identified points or individuals in complex visual environments is to use markers that can be uniquely identified by computer-vision in each picture or frame. Image-based recognition of such markers has been widely used in commercial (e.g. barcodes and Quick-Response, or QR codes) as well as in augmented reality (ARTag, Fiala, 2005) and camera-calibration (CALTag, Atcheson et al., 2010) applications. While such marker-systems have previously been used for high-throughput behavioral studies in ants (Mersch et al., 2013), previous software packages are either not optimized for recognizing isolated tags (as desired for most applications in animal movement), or suffer from licensing issues, making access to these techniques limited. Here, we present and characterize BEEtag (**BE**havioral **E**cology **t**ag), a new open-source software package in Matlab for tracking uniquely identifiable visual markers. First, we provide a basic description of the software and characterize its performance. Next, we validate the tracking system by marking and tracking individual bumblebees (*Bombus impatiens*) within a hive. Finally, we consider the potential extensions, future applications, and limitations of this tracking technique.

## Tag Design and Tracking Software

### Tag design

We use a tag design that is inspired by similar markers for visual tracking such as ARtag (Fiala, 2005) and CALTag (Atcheson et al., 2010). Our tags consist of a 25 bit (5×5) code matrix of black and white pixels that is unique to each tag surrounded by (1) a white pixel border and (2) a black pixel border (Figure 1). The 25-bit matrix consists of a 15-bit identity code, and a 10-bit error check. The 15-bit identity is the binary representation of a number between 1 and 32767, left-padded with zeros and reoriented into a 5×3 pixel matrix (Figure 1A). A unique 10-bit error check is then generated for each code. The first 3 bits of this error code are parity checks (1 (white) for odd and 0 (black) for even) of each of the three columns of the 5×3 code matrix. The next two bits are generated by checking the parity of the first 3 and last 2 columns of the 5×3 code matrix, respectively. This 5-bit error check is then repeated and reversed to give a complete 10-bit error check (Figure 1). This simple binary image matrix can then be scaled to any size where it can be visualized by a camera, for example small tags for use with bumblebees (*Bombus impatiens*, Figure 1B, see below) or moderately larger tags for bigger invertebrates (*Blaberus discoidalis*, Figure 1C, tags roughly 8 mm per side).

**Figure 1.**
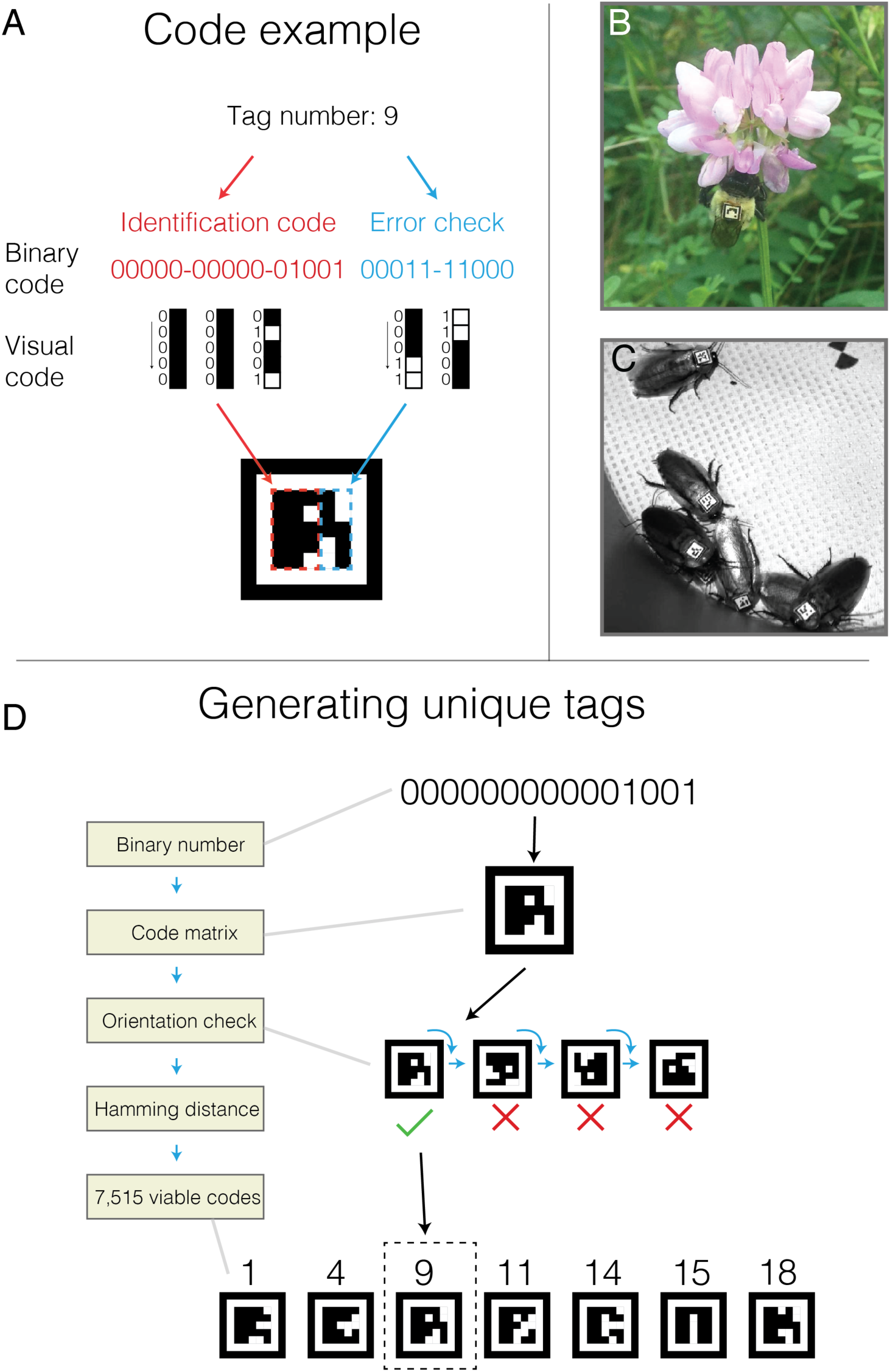
BEEtag code tructure and generation. (A) Basic tag design (see text for details). (B) A bumblebee worker (*Bombus impatiens*) outfitted with a BEEtag and encountered opportunistically in the natural environment. (C) Cockroaches (*Blaberus discoidalis*) outfitted with BEEtags. (D). Schematic representation of the process for generating a list of unique, usable BEEtags.

### Generating a usable set of BEEtags

While a 15 bit encoding theoretically allows for 32,768 different possible code combinations, not all of these can be safely distinguished in practice when the orientation of the tag is unknown (as is the case in most tracking applications). We therefore restrict codes to be used in tracking based on two additional criteria. First, a tag must be valid in only one orientation (i.e. the 10-bit error check matches the 15-bit code in only one of the four possible tag orientations, Figure 1D). Second, any tag must have a Hamming distance of at least 3 (i.e. 3 bits are different) between itself and any valid tag (and its associated alternative orientation). These restrictions, which reduce the number of false positive tag identifications from an image, result in a set of 7,515 viable tags out of the 32,767 possibilities (Figure 1D).

### Identifying BEEtags from an image or video frame

Using this technique, each tag can be uniquely identified in a still image or movie frame without prior knowledge of its position. The raw input for tracking is an image, in color or grayscale format. If tracking tags in a movie, each frame is extracted and analyzed as a separate image. If the frame or still image is in color, it is first converted to grayscale before further processing.

From the grayscale image, the first step is to threshold into a black and white image (Figure 2). In brief, this thresholding step works by converting the matrix of continuous pixel intensity values of an image (i.e. a grayscale image) into a binary matrix using a specified threshold value. This results in a binary (i.e. black and white) image, where zeros are represented by black and ones are represented by white. After converting to a binary image, the software finds all unique regions of white in theimage and checks to see which are rectangular, and all of these regions are considered possible tags (Figure 1C). To verify which regions are true tags and identify them, the software then reads pixel values from within each white rectangle, converts them from black and white values to binary numbers, and references them against the list of viable tags described above. Finally, the position, identity, and orientation of all these tags are recorded and returned to the user as a Matlab structure array.

**Figure 2.**
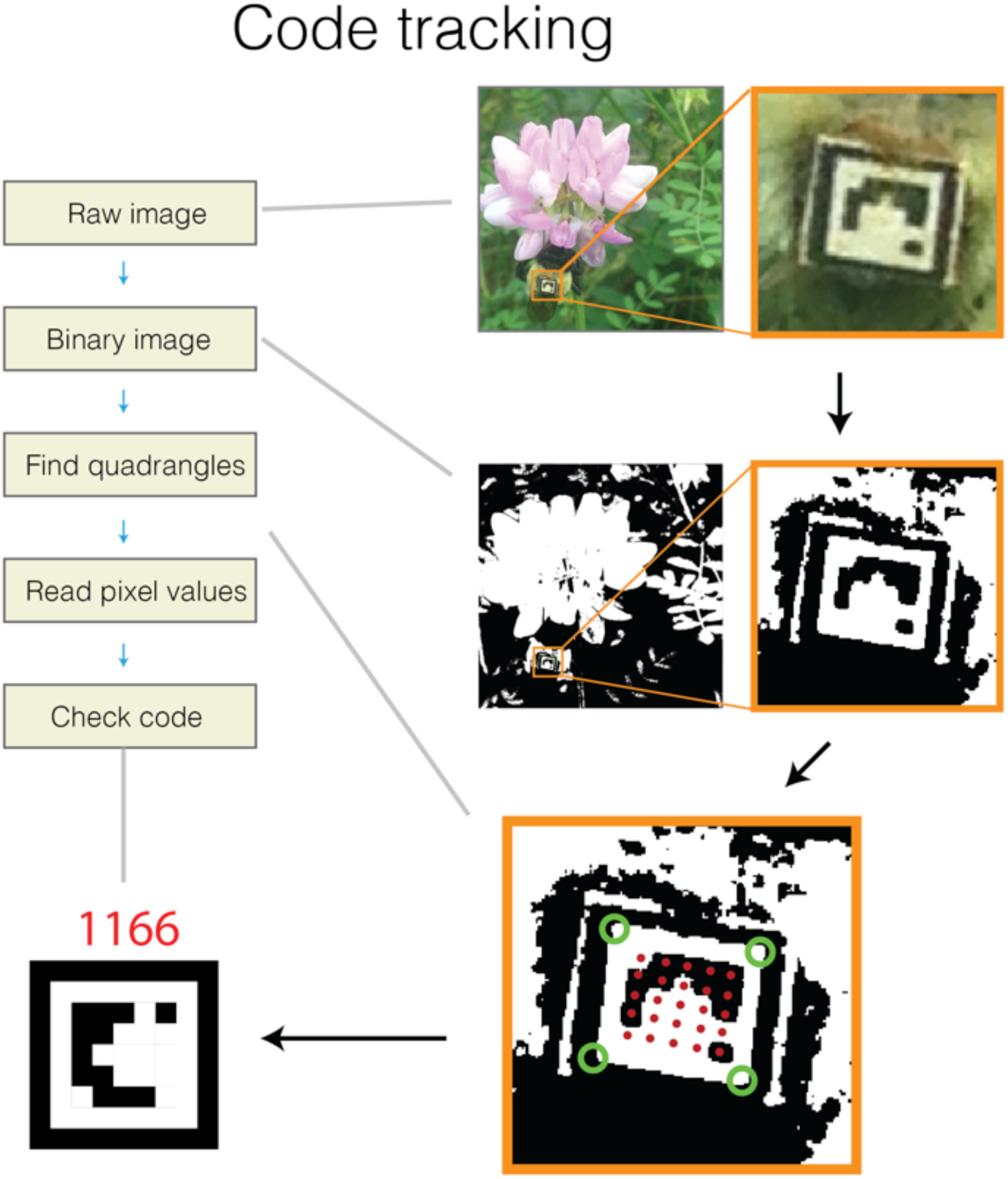
Schematic representation of the algorithm for identify unique BEEtags from an image.Green circles show identified corners of the white quadrangle, and red dots show points where separate pixel values were measured. See text for details.

### Software performance

To test the basic performance of the BEEtag software, we took a video of 12 printed tags with the built-in camera of an iPhone 5 from a constantly moving perspective (Figure 2A, Supplementary Movie 1). We identified codes in each frame while independently varying image resolution, noise level, and black-white threshold levels to examine the effects of these parameters on tracking performance.

In general, tracking performance is strongly affected by all three of these parameters. Resolution was artificially modified using the “imresize” function in Matlab to a range of image resolutions. The average area (in pixels) of the 12 tags in the image was then calculated and the square root of this value taken to estimate the functional resolution of each tag, expressed as the mean length of each tag side (measured as the distance between 2 adjacent corners of the white rectangle containing the tag, Figure 3B). The portion of tags correctly tracked across 255 frames from this sample video dropped dramatically below a resolution of around 25 pixels per tag edge (Figure 3B).

We explored the extent to which noise impairs tracking performance (Figure 3C) by introducing Gaussian noise to each of 100 frames from the sample video using the “imnoise” function in Matlab. This function allowed us to apply Gaussian noise with varying levels of intensity (normalized to an image intensity of 0 to 1) to a full resolution image (i.e. around 38 pixels per tag edge). As expected, increased noise progressively impaired tracking performance, until values of around 0.05 (i.e. variance of 5% of the intensity range) when very few tags were successfully tracked (Figure 2C). Noise impairs tracking by both reducing the efficiency of quadrant tracking and increasing noise within the tag itself. In real applications, noise (i.e. “graininess”) appears in images as a result of unwanted electronic signal, and can depend heavily on the sensor, camera, and recording settings used. For example, digital image noise increases significantly at higher ISO (or light sensitivity of the camera’s sensor) values. In general, however, the noise values reported here are very high (the “0” noise value here represents the direct output of the camera, including digital noise), demonstrating that this tracking system is relatively robust to moderate noise levels. Nevertheless, noise remains an important consideration when designing an image-recording setup.

Finally, black-white thresholding values significantly affected tracking performance (Figure 3D). In parallel to the noise test, we tested the impact of threshold value on tracking performance across 100 frames of the same sample video described above, varying the threshold value over a range from 0.2 to 0.9, corresponding to a normalized intensity value. Tracking performance was optimal at intermediate threshold values, but significantly deteriorated at both low and high threshold values (Figure 3D). Since lighting conditions will vary substantially among real tracking applications, ideal threshold values will vary accordingly (see Validation Experiment below), and therefore finding an optimal tracking threshold will be an important step in each specific application of BEEtag.

**Figure 3.**
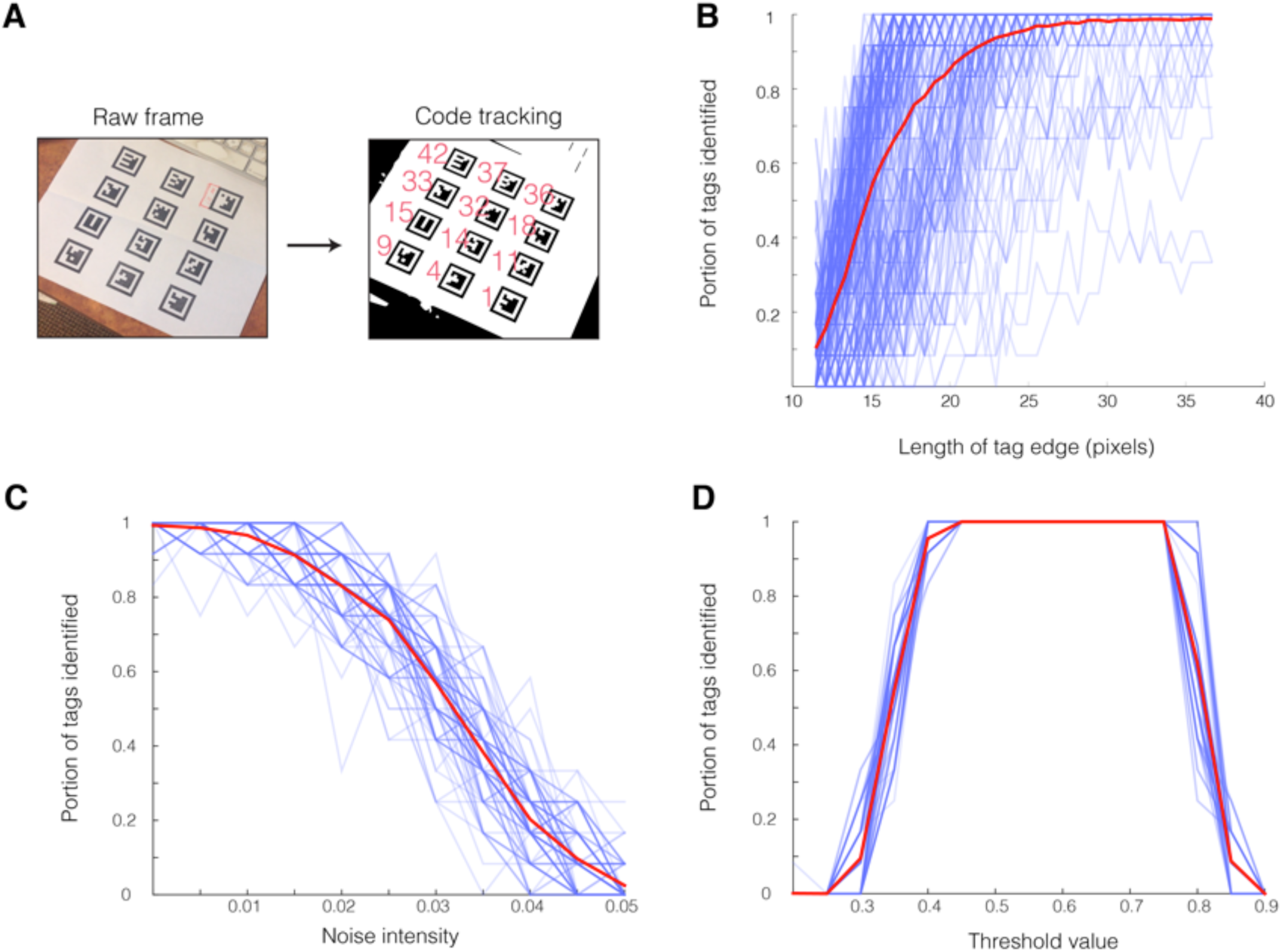
Performance of the BEEtag tracking system in a sample video (A) in response to variation in resolution (B), gaussian noise (C), and binary threshold value (D). See text for details. Transparent blue lines show data from a single video frame (N = 277 in B and N = 100 in C-D), and thickened red lines show the mean across all frames.

Overall, the rate of false positives for tag identification (i.e. the number of tags that are incorrectly identified, rather than not being identified) was low. Among 11,166 codes identified across the combination of 100 images and 15 resolution values described in the resolution test above, 5 were not values actually contained within the image, giving a false positive rate of ∼0.04 % (i.e. 99.96% of codes returned were correctly identified).

### User interface

The BEEtag package consists of a small library of functions available in the supplementary material of this paper. Additionally, a continuously updated respository of the code is available at https://github.com/jamescrall/BEEtag. After installation (i.e. downloading all relevant functions and adding these to the Matlab search path), users interface with the software package primarily through the “locateCodes” function. This function takes a grayscale or color (rgb) image and returns the locations and relevant information (identity, orientation, etc) of any BEEtag tags located in the image as a Matlab structure array. Users have the option of manually specifying the threshold value for binary image conversion, size limits for tags in pixels, and other visualization options.

## Experimental validation: Spatial behavior patterns in a bumblebee hive

### Study species and tag attachment

To validate the BEEtag tracking system, we outfitted individual bumblebees (*Bombus impatiens*) with unique BEEtags to track spatial movement of multiple individuals simultaneously within the hive. A single hive (Biobest) was maintained indoors but with access to the natural environment through a plastic tube, which allowed the bees to enter and exit the hive at will to forage for nectar and pollen. The hive was initially placed on July 9th and given seven days to acclimate and begin normal foraging activity. On July 16th, we outfitted roughly 100 workers with unique BEEtags. All BEEtags used were printed on a single 8.5 × 11”sheet of waterproof, tear-resistant paper on a high-resolution (1200 dpi) laserjet printer at Staples®. Each tag was cut out from the sheet by hand, measured roughly 2.1mm × 2.1 mm, and weighed around 1.83 mg. All bees except the queen were removed from the hive at the same time using a vacuum aspirator (Bioquip Products) and maintained for 30-60 min at 4° C to reduce activity level. Bees were then individually cold-anaesthetized at -20° C and outfitted with a unique tag attached with cyanoacrylate gel glue. After tagging, all bees were then returned to the hive and allowed to acclimate for 24 hours before data collection and data collection occurred on July 17th.

### Imaging setup and data collection

To capture images of sufficiently high resolution to track individual tags over the entire hive arena (roughly 21.5 × 15.0 cm), we used an entry-level DSLR camera (Nikon D3200), operating at the maximum resolution of 6016 × 4000 pixels per image. The hive was outfitted with a clear plexiglass top prior to data collection and illuminated by three red lights, to which bees have poor sensitivity (Briscoe and Chittka, 2001). The camera was placed ∼ 1 m above the hive top and triggered automatically with a mechanical lever driven by an Arduino microcontroller. On July 17th, pictures were taken every 5 seconds between 12:00 pm and 12:30 pm, for a total of 372 photos. 20 of these photos were analyzed with 30 different threshold values to find the optimal threshold for tracking BEEtags (Figure 4M), which was then used to track the position of individual tags in each of the 372 frames.

**Figure 4.**
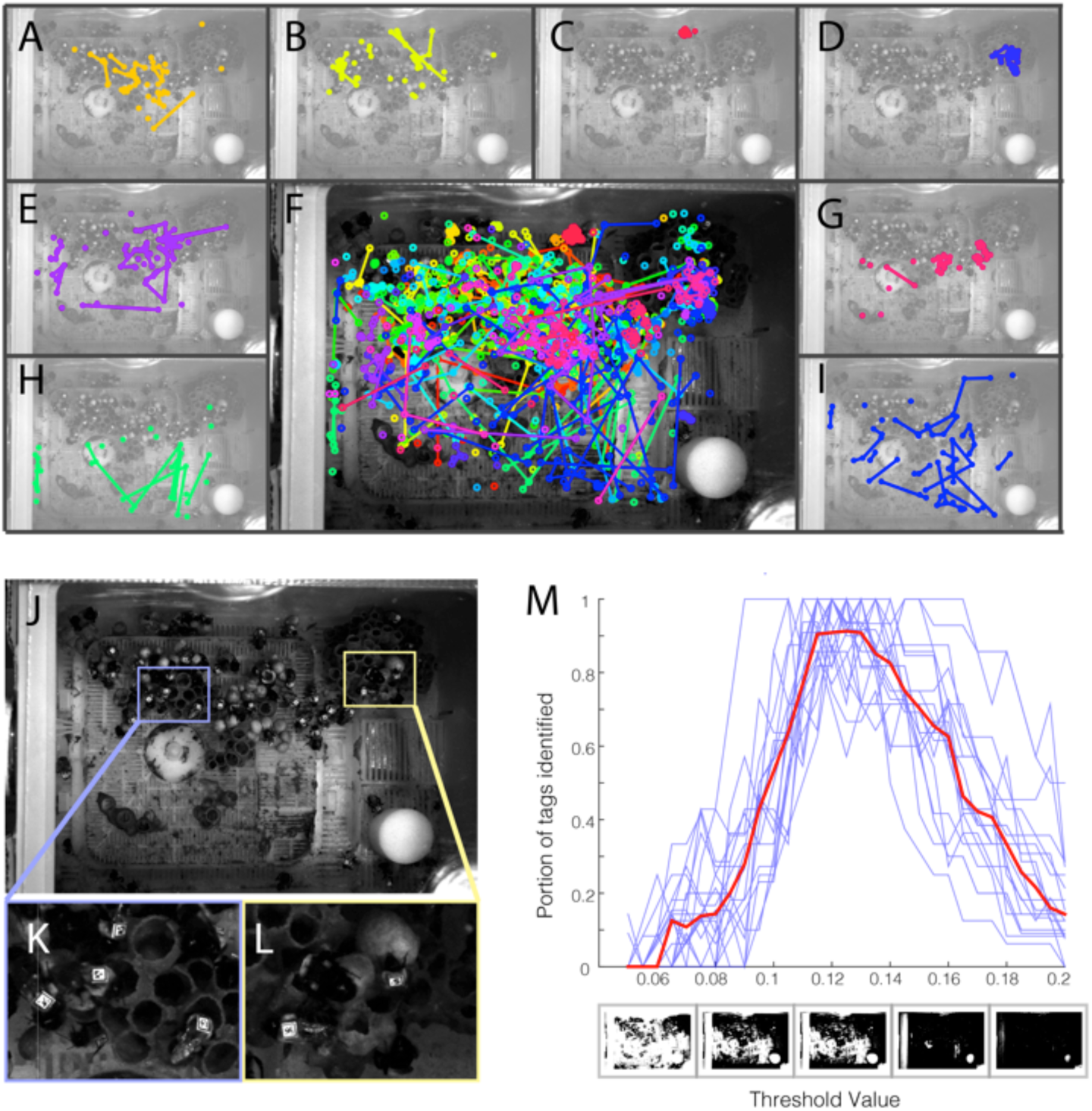
Validation of the BEEtag system in bumblebees (*Bombus impatiens*). (A-E, G-I) Spatial position over time for 8 individual bees, and (F) for all identified bees at the same time. Colors show the tracks of individual bees, and lines connect points where bees were identified in subsequent frames. (J) A sample raw image and (K-L) inlays demonstrating the complex background in the bumblebee hive. (M) Portion of tags identified vs. threshold value for individual pictures (blue lines) and averaged across all pictures (red line).

### Results and tracking performance

Overall, 3516 locations of 74 different tags were returned at the optimal threshold. In the absence of a feasible system for verification against human tracking, false positive rate can be estimated using the known range of valid tags in the pictures. Identified tags outside of this known range are clearly false positives. Of 3516 identified tags in 372 frames, one tag (identified once) fell out of this range and was thus a clear false positive. Since this estimate does not register false positives falling within the range of known tags, however, this number of false positives was then scaled proportionally to the number of tags falling outside the valid range, resulting in an overall correct identification rate of 99.97%, or a false positive rate of 0.03%.

Data from across 30 threshold values described above were used to estimate the number of recoverable tags in each frame (i.e. the total number of tags identified across all threshold values) estimated at a given threshold value. The optimal tracking threshold returned an average of around 90% of the recoverable tags in each frame (Figure 4M). Since the resolution of these tags (∼33 pixels per edge) was above the obvious size threshold for optimal tracking (Figure 3B), untracked tags most likely result from heterogeneous lighting environment. In applications where it is important to track each tag in each frame, this tracking rate could be pushed closer to 100% by either (a) improving lighting homogeneity or (b) tracking each frame at multiple thresholds (at the cost of increased computation time).

These locations allow for the tracking of individual-level spatial behavior in the hive (see Figure 4F) and reveal individual variations in both activity and spatial preferences. For example, some bees remain in a relatively restricted portion of the hive (e.g. Figure 4C and D) while others roamed widely within the hive box (e.g. Figure 4I). Spatially, some bees restricted movement largely to the hive pots and developing brood (e.g. Figure 4B), while others tended to remain off the hive pots (e.g. Figure 4H) or showed mixed spatial behavior (e.g. Figure 4A, E, and G).

## Discussion

Here, we have presented a new open-source software package – BEEtag - for tracking unique visual markers and demonstrated its utility for studies of animal behavior. This package builds directly off previous work aimed at tracking individually identifiable markers (Atcheson et al., 2010; Fiala, 2005) and extends previous efforts by providing a simple interface in Matlab that is intended to improve ease of use for researchers in behavioral ecology and other branches of the life sciences.

Tracking systems that utilize uniquely identifiable markers such as BEEtag (or ARTag and CALTag) have some fundamental advantages over other techniques. One primary advantage is that tags are identified independently in each photo or frame, so errors don’t propagate across frames. For example, in most automated tracking systems (e.g. (Branson et al., 2009; de Chaumont et al., 2012; Hedrick, 2008), with notable exceptions such as (Pérez-Escudero et al., 2014)), individual tracking depends on information from previous frames, and therefore when an individual is either (a) not tracked or (b) incorrectly tracked in one or a few frames (i.e. because the individual is occluded from view or interacts with another individual), tracking fails (Pérez-Escudero et al., 2014). While acceptable for short-term data-collection, longer-term longitudinal data sets (as are often particularly relevant for behavioral ecology) are difficult or impossible to collect with such techniques.

Another important advantage of this tracking system is that it does not require a homogenous background, as do many optical tracking systems (Branson et al., 2009; de Chaumont et al., 2012; Pérez-Escudero et al., 2014). While it is possible in a controlled laboratory setting to create a homogenous background for automated detection of image regions associated with an animal’s body, this is difficult or impossible in most naturalistic contexts (Dell et al., 2014). BEEtags, on the other hand, are robust to complexity in the background image (see Figure 1B and Figure 4J-L [although not necessarily lighting heterogeneity, Figure 4M, see discussion above]). For example, the sample image used in Figure 2 of a bumblebee worker with a BEEtag was taken opportunistically with an iPhone 5 against a natural background when the bee was encountered foraging outside of the hive, and emphasizes the robustness of this tracking system in natural environments.

Another important advantage of the BEEtag system is its cost. The examples included here used either an iPhone 5, or a commercially available Nikon DSLR camera (currently available for ∼$500 USD), and tags were printed on waterproof, tear-resistant paper at a cost of $0.87 USD for 600 tags (approximately 0.145 cents each). This system thus makes the collection of high-quality, high-throughput behavioral datasets possible at an extremely low cost compared to alternative systems.

Like all other tracking systems, however, BEEtag has limitations that make it better suited to certain applications than others. First, the system requires the application of a tag. Handling (Pankiw and Page, 2003) and tag application (Dennis et al., 2008) can significantly affect stress levels (Sockman and Schwabl, 2001) and behavior in animals (Ropert-Coudert and Wilson, 2005). While BEEtags are lightweight (depending on size and printing material), the potential biomechanical and behavioral effects of both tag attachment (Aldridge and Brigham, 1988) and handling need to be carefully considered for each study organism and specific application.

Since BEEtag depends on visual information, performance also can be substantially affected by (a) uneven lighting (see above and Figure 4M), (b) animal posture, and (c) tag cleanliness. While issues of uneven lighting can be computationally overcome by either identifying codes at multiple threshold values or applying an appropriate high-pass spatial filter to images, the other limitations are more fundamental and mean that BEEtag tracking performance will be impaired in situations where tags are not visible (i.e. when animals are piled on top of each other) or cannot be kept clean (potentially an important consideration for freely behaving animals in natural environments).

Another important limitation when considering the utility of BEEtag for specific applications are the challenges of data storage and processing, which can be significant for any image processing techniques when compared to alternative tracking techniques such as RFID (Henry et al., 2012). While performing processing in real time can minimize data storage problems, this is not possible in all applications. In particular, large images such as those used in the validation experiment described above (Figure 4) can be computationally intensive, and therefore impractical for real-time processing.

### Alternative application and future directions

While we have focused here on using BEEtags for tracking the overall spatial position of individuals, the utility of this tracking system is not limited to ethology or behavioral ecology. One such potential direction that seems particularly promising is use in the field of animal locomotion. Focus in the study of animal locomotion has increasingly shifted from steady-state locomotion in laboratory environments to dynamic movement in complex, naturalistic environments (Combes et al., 2012; Daley and Biewener, 2006; Dickinson et al., 2000), where tracking is particularly challenging (Dell et al., 2014). Since having tags obscured for some or many frames is not highly problematic for BEEtag, we suggest that this tagging system could be of particular utility for tracking the kinematics of animal locomotion through cluttered environments, where they are likely to be temporarily obscured. Additionally, in applications where multiple rigid points are tracked in order to, for example, reconstruct three-dimensional body rotations (Ravi et al., 2013), these points could be automatically extracted from a properly oriented BEEtag, thereby negating the need for manual or semi-automated digitizing (Hedrick, 2008).

The BEEtag package will be maintained regularly on the GitHub site, which allows for user contributions, and it is our hope that as use of this software increases, users will contribute improvements, modifications, and extensions that will both improve performance and ease of use to the current implementation of BEEtag, as well as extending this technology to new applications.

## Acknowledgements

We are grateful to Benjamin de Bivort, Dominic Akandwanaho, Sawyer Hescock, and Andre Souffrant for help in debugging tracking software and placement of BEEtags on cockroaches. This work was funded by an NSF GRFP fellowship to James Crall and an NSF CAREER grant to Stacey Combes (IOS-1253677). Nick Gravish would like to acknowledge funding from the James S. McDonnell foundation.

